# Heat shock induces alternative polyadenylation through dynamic DNA methylation-regulated chromatin looping

**DOI:** 10.1101/2023.08.25.554792

**Authors:** Emily E. Fink, Vishal Nanavaty, Byron H. Lee, Angela H. Ting

## Abstract

Alternative cleavage and polyadenylation (APA) is a gene regulatory mechanism used by cells under stress to upregulate proteostasis-promoting transcripts, but how cells achieve this remains poorly understood. Previously, we elucidated a DNA methylation-regulated APA mechanism, in which gene body DNA methylation enhances distal poly(A) isoform expression by blocking CTCF binding and chromatin loop formation at APA control regions. We hypothesized that DNA methylation-regulated APA is one mechanism cells employ to induce proteostasis-promoting poly(A) isoforms. At the *DNAJB6* co-chaperone gene locus, acute heat shock resulted in binding of stress response transcription factors HSF1, ATF6, and YY1 at the APA control region and an increase in the expression of the proximal poly(A) isoform known to prevent protein aggregation. Furthermore, TET1 was recruited to rapidly demethylate DNA, facilitating CTCF binding and chromatin loop formation, thereby reinforcing preferential proximal poly(A) isoform expression. As cells recovered, the transcription factors vacated the APA control region, and DNMT1 was recruited to remethylate the region. This process resolved chromatin looping and reset the poly(A) isoform expression pattern. Our findings unveil an epigenetic mechanism enabling cells to dynamically modulate poly(A) isoforms in response to stress while shedding light on the interplay between DNA methylation, transcription factors, and chromatin looping.

## INTRODUCTION

The cellular response to perturbations in proteostasis includes the compartment-specific heat shock response (HSR) in the cytosol and the unfolded protein response (UPR) in the endoplasmic reticulum (ER), with significant crosstalk between both pathways (1). The HSR and UPR are multifaceted and evolutionarily conserved signalling pathways aimed at inducing an anti-stress proteome. If the cell is unable to resolve the stress, apoptosis ensues (2). In eukaryotic organisms, the cascade of signals involved in the cell stress response has been significantly studied at the transcriptional, translational, and post-translational levels. The canonical events involved in the HSR include the induction of the transcription factor Heat Shock Factor 1 (HSF1), which promotes the transcription of heat shock proteins (3). The key events involved in the UPR include the post-translational activation of ER-membrane associated proteins, IRE-1, PERK, and ATF6, upon presence of misfolded proteins in the ER lumen, followed by translational attenuation, regulated IRE1-dependent decay of mRNA (RIDD), and selective transcriptional upregulation of proteostasis-promoting genes via transcription factors XBP-1s, ATF4, and ATF6 (1,2,4-6). Furthermore, recent efforts have highlighted additional layers of regulation during the cell stress response at the level of post-transcriptional mRNA processing, specifically alternative cleavage and polyadenylation (APA) (5,7).

About 70-80% of human genes can be terminated at multiple polyadenylation [poly(A)] sites, giving rise to distinct mRNA isoforms with contrasting properties. These alternative poly(A) isoforms can differ in their 3’ untranslated region (UTR) lengths or in amino acid coding content. Differences in 3’ UTR lengths can affect mRNA metabolism and ultimately the expression of the final protein. For example, longer 3’ UTRs can harbour binding sites for RNA binding protein or microRNA, which can modulate mRNA half-life, alter mRNA localization, or modify translation efficiency (5). Alternative intronic poly(A) sites can generate distinct protein isoforms, many of which have unique functions. Thus, it is not surprising that the cellular response to proteotoxic stress can utilize APA as a mechanism to selectively express proteostasis-promoting transcripts while simultaneously suppressing overall transcription and translation. For instance, 3’ UTR global shortening has been observed in cells exposed to arsenic stress as an adaptive mechanism to preserve mRNAs during recovery from stress (5). Global APA changes have also been observed following stressors such as heat shock (8), nutrient deficiency (9), DNA damage (10), and oxidative stress (5,11). While the underlying mechanism dictating poly(A) site usage is largely unknown, it has been suggested that epigenetic features and chromatin structure can be altered during stress and recovery (12) and should be explored as a potential mechanism.

DNA methylation changes have been observed during environmental heat shock of organisms within the livestock industry (13,14). Pathophysiological hypermethylation of ER stress genes has been observed in association with insulin resistance (15,16) and cancer (17). DNA demethylation has also been observed during the cellular response to ER stress in lens epithelial cells (18). While stress-induced alterations in DNA methylation patterns have been documented, their impact on the transcriptome is well understood. Our group recently described a link between DNA methylation, chromatin structure, and APA (19), where CCCTC-binding factor (CTCF) can bind to DNA demethylated regions between poly(A) sites in a gene to facilitate CTCF/cohesin-mediated chromatin looping, which leads to the physical blockade of RNA polymerase II and subsequent preferential expression of proximal poly(A) isoforms. Here, we postulate that DNA methylation could be modulated during cellular stress response as a potential mechanism upstream of APA, driving the expression of proteostasis-promoting transcripts through CTCF-mediated chromatin loop formation.

To test this hypothesis, we focused on the co-chaperone gene DnaJ Heat Shock Protein Family (Hsp40) Member B6 (*DNAJB6)*, previously identified by our group as a target of DNA methylation-regulated APA (19). Following exposure to heat-shock, we assessed the temporal changes in DNA methylation, *DNAJB6* poly(A) isoform expression, chromatin loop formation, and protein expression. We observed dynamic changes in DNA methylation during heat shock and recovery at the putative *DNAJB6* APA control region. The DNA methylation shifts correlated with *DNAJB6* mRNA and protein isoform expression. In addition, we assessed the temporal recruitment of several DNA binding factors and DNA methylation machinery at this locus in the context of heat shock to begin unravelling the mechanism for dynamic DNA methylation remodelling during stress response. Altogether, our data illustrates a surprising mechanism connecting the dynamic DNA methylation landscape to chromatin looping and ultimately poly(A) isoform expression during the cellular response to heat shock.

## MATERIAL AND METHODS

### Cell culture

HCT116 and HEK-293 cells were obtained from the ATCC. DNMT1 and 3b double knock out (DKO) cells were obtained from the laboratory of Dr. Bert Vogelstein (20). HCT116 and DKO were cultured in McCoy’s 5a Medium Modified (Thermo Fisher Scientific) supplemented with fetal bovine serum (FBS) (Corning) to a final concentration of 10%. The HEK-293 cells were cultured in Eagle’s Minimum Essential Medium (Thermo Fisher Scientific) supplemented with FBS (Corning) to a final concentration of 10%. All cells were kept in an incubator at 37□°C in with 5% CO_2_. For heat shock, cells were incubated at 43°C for one hour. Cells were harvested immediately after the one-hour incubation for the time 0 time point and at 1, 2, 4, 12, 24, and 48 hours after recovery at 37°C. DAC treatment was performed as described previously (19).

### dCas-directed DNA demethylation

To establish HCT116 cell lines that expressed the dCas9-TET demethylation system as described previously (21). First, lentivirus containing either control or the DNAJB6 gRNA expression construct (VectorBuilder) was made using HEK-293 cells, following the standard LipoD293 transfection protocol (SignaGen Laboratories), with the envelope (pCMV-VSV-G, AddGene) and packaging (psPAX2, AddGene) plasmids in a 2.5:1 ratio. At 48 hours post-transfection, lentivirus-containing media was collected from the HEK-293 cells, filtered, and added to the HCT116 cells in the presence of 8 μg/mL polybrene (Sigma-Aldrich). Lentivirus-containing media was replaced after 24 hours with regular culture media supplemented with 10% FBS, and infected cells were selected using 1 μg/ml puromycin (Sigma-Aldrich) for 48 hours. Following selection, the HCT116 cells were transfected with both pdCas9-Tet1-CD (Addgene) and pcDNA3.1-MS2-Tet1-CD (Addgene) plasmids using Lipofectamine 2000 (Life Technologies) according to the manufacturer’s instructions. Cells were harvested 2 days post transfection for RNA and genomic DNA extraction.

### Chromatin Immunoprecipitation (ChIP)

ChIP was performed using the Magna ChIP A/G kit (Sigma-Aldrich) following manufacturer’s protocol. The antibodies and their respective dilutions are as follows: CTCF antibody (Cell Signaling Technology #2899) at 1:25 dilution, YY1 antibody (Cell Signaling Technology #46395) at 1:50 dilution, ATF6 antibody (Cell Signaling Technology #65880) at 1:50 dilution, TET1 antibody (ThermoFisher #61443) at 1:25 dilution, HSF1 antibody (Novus #NBP2- 42206V) at 1:50 dilution, and DNMT1 antibody (Sigma-Aldrich #D4692) at 1:50 dilution.

### DNA pulldown

The DNA pulldown protocol was adapted from the previously published procedure (22) with a few minor changes. DNA probes (chr7:157,204,244-157,204,652, GRCh37/hg19) spanning the APA control region in *DNAJB6* were generated using PCR (EpiMark-HS-TAQ, New England Biolabs), where one set of reactions used a 5-methylcytosine-containing dNTP mix (methylated probes) and the other set used the dNTP mix without methylated bases (unmethylated probes). Terminal transferase (New England biolabs) was used to add a biotinylated dUTP tail. DNA probes were incubated with nuclear lysates from untreated HCT116 cells. DNA/protein complexes were extracted using streptavidin magnetic beads, and a bead only control was included as a third reaction. Eluted proteins from each reaction were resolved using SDS-PAGE. The DNA pulldown experiment was performed in biological duplicates.

The gel was stained with Coomassie Blue and subjected to in-gel protein digestion for LC-MS sample preparation. Briefly, each lane in the gel was divided into smaller pieces, washed (50% ethanol), de-stained (5% acetic acid), dehydrated (acetonitrile), dried, reduced, alkylated, and then digested with trypsin (5 µL of 10 ng/µL trypsin in 50 mM HCO_3_) overnight at room temperature. The peptides were extracted from polyacrylamide with two aliquots of 30 µL 50% acetonitrile with 5% formic acid. The aliquots were combined and evaporated to <10 µL in Speedvac and then resuspended in 1% acetic acid to a final volume of 30 µL for LC-MS analysis using a ThermoScientific Fusion Lumos mass spectrometry system. The data were analysed by using all CID spectra collected in the experiment to search against the human SwissProt database and the sequences of Human HIF-2-alpha (accession Q99814) using the program Sequest. The search results were further validated with Scaffold (Scaffold_4.7.2, Proteome Software Inc.).

Within each biological replicate, proteins with less than 10 spectral counts were filtered out. Next, only those proteins with a ≥ 5-fold enrichment of spectral counts over bead control were considered positive. Finally, 139 proteins were reproducibly identified in both biological replicates (**Supplemental Table 2**) and further annotated as chromatin binding (GO:0003682), transcription factor binding (GO:0008134), DNA-binding transcription factor binding (GO;0140297), sequence-specific DNA binding (GO:0043565), DNA secondary structure binding (GO:0000217), or transcription regulator activity (GO:0140110) using g:Profiler (23).

### Targeted bisulfite sequencing

DNA was bisulfite converted using the EZ DNA methylation-gold kit (Zymo #D5006). *DNAJB6* loci of interest were amplified using the primers listed in **Supplemental Table 3**. Amplicons were resolved on 1% agarose gel by gel electrophoresis and purified using the QiaQuick Gel Extraction Kit (QIAGEN #28706). Purified PCR product was cloned into the TOPO TA Vector and transformed into OneShot TopTen Chemically Competent *E. coli* cells (Thermo Fisher Scientific #K457540). Alleles were analysed using BISMA (24) after Sanger sequencing.

### Quantification of poly(A) isoform expression by qRT-PCR

Total RNA was extracted by AllPrep kit (Qiagen). Oligo-dT(16) primer was used to convert 1 μg of total RNA into cDNA using Superscript III reverse transcriptase (Thermo Fisher #18080093), and 40 ng of cDNA was used for each qPCR reaction with QuantiTect SYBR green reagent (QIAGEN #204143). PCR primers for specific poly(A) isoforms are listed in **Supplemental Table 3**. Relative isoform expression for *DNAJB6* was calculated as distal isoform level divided by proximal isoform level. Results were calculated from biological triplicates that were assayed by technical triplicate PCR reactions, and statistical significance was determined using ordinary one-way ANOVA with multiple comparisons correction in GraphPad Prism 9.0.0.

### Chromatin conformation capture (3C)

3C was performed following previously optimized protocols (19,25) with minor changes. In brief, a total of 10^7^ cells were harvested with trypsin, washed, and re-suspended in 9.5 mL PBS. The samples were cross-linked in 1% formaldehyde for 10 minutes at room temperature with rotation. Nuclei were extracted by incubating cells on ice for 10 minutes using 5 mL cold lysis buffer [10 mM Tris-HCl, pH 7.5; 10 mM NaCl; 5mM MgCl_2_; 0.1 mM EGTA; 1X complete protease inhibitor (Roche #11836145001)]. Pelleted nuclei were re-suspended in 0.5 mL 1.2x restriction enzyme buffer with 0.3% SDS and incubated for 1 hour at 37°C while shaking at 900 rpm, followed by the addition of Triton X-100 to 2% final concentration and incubated for 1 hour at 37°C while shaking. 400 U of BamHI (NEB #R1036S) was added, and cells were incubated overnight at 37°C with shaking. SDS was added to 1.6% final concentration, and samples were incubated at 65°C for 20 minutes with shaking. Ligation of the digested DNA was carried out by adding 6.125 mL of 1.15x ligation buffer (10X: 660 mM Tris-HCl, pH 7.5; 50 mM DTT, 50 mM MgCl_2_, 10 mM ATP) and adding Triton X-100 to a final concentration of 1%, followed by incubation for 1 hour at 37°C with gentle shaking. Samples were incubated with 100 U of T4 DNA ligase (NEB #M0202S) for 4 hours at 16°C followed by 30 minutes at room temperature. Samples were then de-crosslinked overnight at 65°C with 300 μg proteinase K (Invitrogen #25530049) followed by incubation with 300 μg RNase A (Invitrogen #EN0531) for 30 minutes at 37°C. DNA was purified using phenol/chloroform extraction and re-suspended in 150 μl of 10 mM Tris pH 7.5. DNA was quantified using one primer adjacent to the BamHI cut site in each distal fragment and another one within the anchor region (**Supplemental Table 3**) by SYBR green real-time quantitative PCR on the Roche LightCycler® 96 on 50X diluted 3C DNA and serial dilutions of reference DNA of known concentration using internal primer sets that do not amplify across BamHI cut sites.

### Western blot

Cells were lysed in 3% SDS in 10mM Tris pH = 7.5 by pipetting then centrifuge through a Qiashredder column (Qiagen #79656). Protein concentrations were determined by BCA assay (Thermo Fisher Scientific #23225). 10 μg of cell lysate per sample were resolved on SDS-PAGE in NuPAGE reducing sample buffer (Thermo Fisher Scientific #NP0001) using the Novex system at 120V for 1 hour. Proteins were transferred to PVDF membrane using a 1X transfer buffer (Thermo Fisher Scientific #NP0006-1) with 10% methanol at 30V for 3 hours at 4°C. Membranes were blocked in 10% non-fat dairy milk in 1X TTBS (25mM Tris-HCl, 155 mM NaCl, 0.1% Tween 20) overnight at 4°C. Membranes were incubated either at room temperature for 2 hours or overnight at 4°C with primary antibodies in 5% milk in 1X TTBS at the following concentrations: DNAJB6 (Abcam #ab198995; 1:1000), CTCF (Cell Signaling Technology #2899; 1:1000), YY1 (Cell Signaling Technology #46395; 1:1000), DNMT1 (Sigma-Aldrich #D4692; 1:1000) and TET1 (Thermo Fisher Scientific #61443; 1:500). Rat and mouse secondary antibodies were used at 1:5000 in 5% milk in 1X TTBS with 1.5-hour incubation at room temperature. ACTB primary antibody (Sigma Aldrich #A3584) was used at 1:10,000 dilution in 5% blocking buffer with a 1-hour incubation at room temperature. Membranes were washed five times for 5 minutes each in 1X TTBS at room temperature after primary and secondary antibody incubations. Membranes were exposed to film following incubation with ECL Plus (Thermo Scientific #32132).

## RESULTS AND DISCUSSION

### *DNAJB6* isoform expression is mediated by DNA methylation

We previously described a mechanism by which DNA methylation regulates APA through CTCF/cohesin-mediated chromatin looping (19). We identified 546 genes undergoing APA between a pair of isogenic colon cancer cell lines that differ in their genomic DNA methylation levels. A subset of these genes has differential DNA methylation between their alternative poly(A) sites. In the absence of DNA methylation, CTCF binds its recognition motif in these regions to facilitate cohesin complex docking and chromatin loop contact formation. Coincident with the loop contact point, we also observed recruitment of serine 5 phosphorylated RNA polymerase II (Pol2Ser5) and enrichment of histone H3 lysine 27 acetylation (H3K27Ac), reminiscent of an enhancer-promoter interaction. The large protein cluster consisted of CTCF, cohesin, and PolSer5 presented a roadblock for the transcription elongation complex, which decreased the use of the distal poly(A) site and leading to preferential expression of the proximal poly(A) isoform. Conversely, DNA methylation at these same regions prevents CTCF binding, thus no loop forms between the poly(A) sites, and RNA polymerase II can more readily reach the distal poly(A) site. The extent of the interplay between DNA methylation, three-dimensional chromatin structure, and its impact on APA has yet to be fully explored in eukaryotic cells. To gain insight into potential pathways and physiological states where this mechanism may play an important role, we performed Gene Ontology (GO) analysis on the 546 genes (**Supplemental Table 1**). The enriched terms hinted at cellular proteostasis-related functions and processes. The top enriched Molecular Functions (MF) terms included protein binding, enzyme binding, cadherin binding, and cytoskeletal protein binding. The top enriched Biological Processes (BP) terms were biological regulation, regulation of protein metabolic process, establishment of localization in a cell, regulation in response to stress, and regulation of protein catabolic process. To explore this potential connection between DNA methylation-regulated APA and proteostasis, we decided to focus on one of the candidate genes with a role in protein turnover, DnaJ heat shock protein family (Hsp40) member B6 (*DNAJB6*) (26).

DNAJB6 is a co-chaperone protein that functions together with Hsp70 chaperones to prevent protein aggregation and misfolding. It is a highly conserved and ubiquitously expressed member of the Hsp40 family of proteins. DNAJB6 abnormalities have been associated with several pathologies including Huntington’s disease (26-28), breast cancer (29,30), Alzheimer’s disease (31), limb-girdle muscular dystrophy (32), and other synucleinopathies (33). *DNAJB6* gene is located on chromosome 7q36.3 and has two annotated isoforms in the reference genome (**Figure 1A**). Previously generated poly(A) sequencing data (19) showed preferential usage of the distal poly(A) site in the DNA methylation-competent HCT116 cells but higher usage of the proximal poly(A) site in the methylation-deficient DKO cells. By comparing HCT116 cells to DKO, we identified a CpG island (chr7:157,204,349-157,204,587, GRCh37/hg19) between the two poly(A) sites to harbour differential DNA methylation and CTCF/cohesin binding (**Figure 1B**). The poly(A) isoform expression pattern is consistent with what our model of DNA methylation-regulated APA would predict: in the absence of DNA methylation (DKO cells), CTCF and cohesin bind to the CpG island to promote proximal poly(A) isoform expression; in the presence of DNA methylation (HCT116 cells), the lack of CTCF/cohesin binding enables higher distal poly(A) isoform expression. The two mRNA isoforms are translated into two distinct proteins (**Figure 1C** & **1D**) that share an identical N-terminal J-domain responsible for interacting with the ATPase domain of Hsp70, as well as an identical G/F domain. The C-termini of both isoforms are serine-rich but differ in length, where the distal isoform sequence contains a nuclear localization signal not found in the proximal isoform (**Figure 1D**).

**Figure 1.**
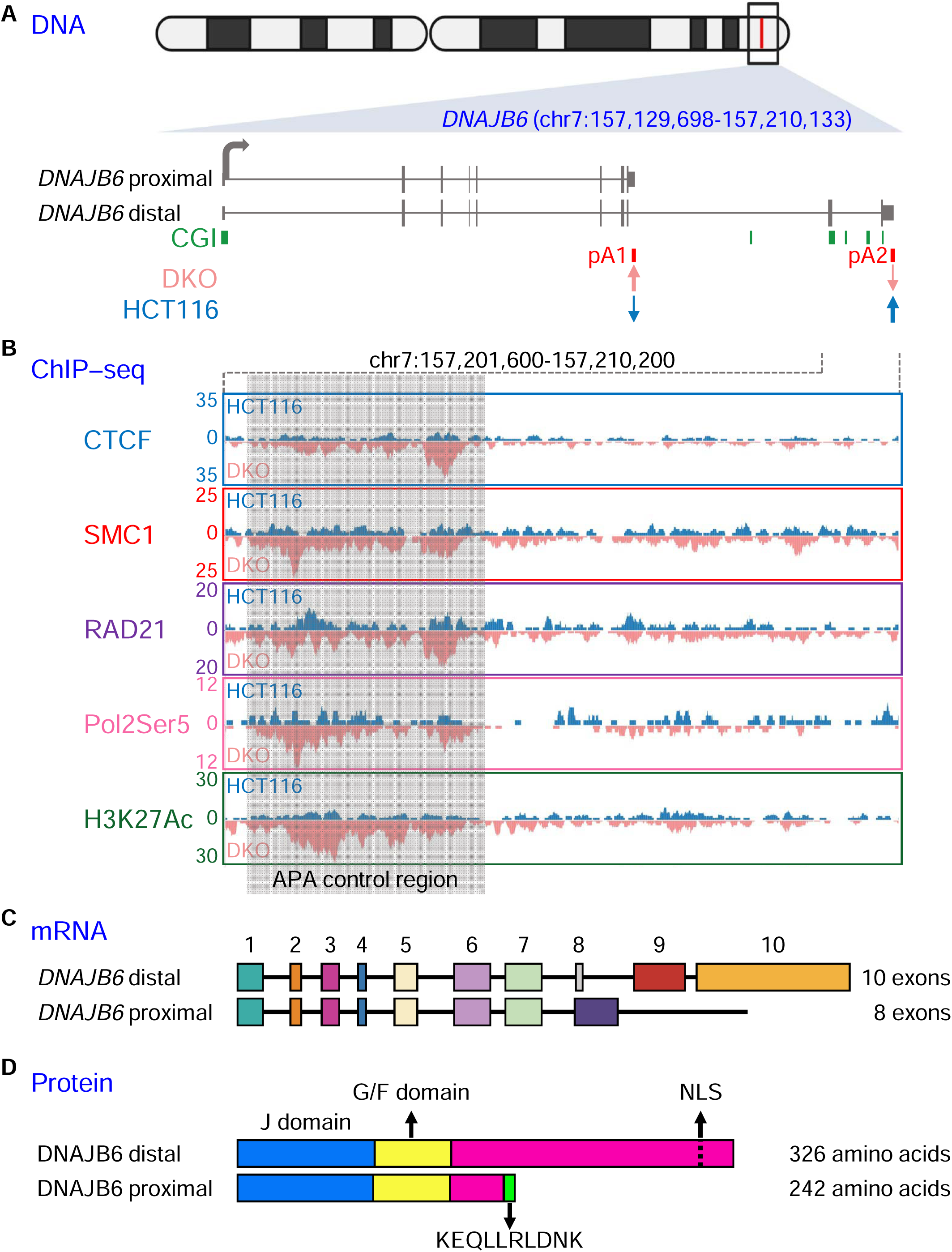
*DNAJB6* is a target of DNA methylation-regulated APA. (**A**) Genomic and poly(A) site locations of *DNAJB6*. DKO cells have higher usage of the proximal poly(A) site (pA1) while HCT116 cells relatively higher usage of the distal poly(A) site (pA2). CGI; annotated CpG islands. (**B**) UCSC genome capture of ChIP-seq data in HCT116 (blue) and DKO (pink) cells for CTCF, SMC1, RAD21, serine 5 phosphorylated RNA polymerase II (Pol2Ser5), and histone H3 lysine 27 acetylation (H3K27Ac) surrounding the putative *DNAJB6* APA control region. (**C**) Schematic of mRNA isoforms. (**D**) Schematic of protein isoforms.

The two isoforms have been reported to have distinct functions, presumably due to their differential cellular localization. While the proximal DNAJB6 isoform is predominantly cytoplasmic, the distal isoform is primarily nuclear and has been implicated as a tumour suppressor in several malignancies through its nuclear activities in modulating the WNT signalling pathway (26,29,34,35). Both isoforms are capable of suppressing protein aggregation in their respective compartments (26,36). As such, the disruption of DNAJB6 activity has been the focus of several diseases characterized by abnormal protein aggregation, including Huntington’s disease (26-28), Parkinson’s diseases (33), limb-girdle muscular dystrophy (32), and Alzheimer’s disease (31). While the expression pattern of DNAJB6 isoforms has been studied in recent years, the mechanism regulating isoform expression is not well understood. Based on our data (**Figure 1B**), we speculate that the differentially methylated CpG island is a regulatory sequence involved in APA modulation of *DNAJB6*.

### Perturbation of DNA methylation at the APA control region alters *DNAJB6* isoform expression

To confirm differential DNA methylation of this putative APA control region between HCT116 and DKO cells, we preformed targeted bisulfite sequencing across this CpG island. HCT116 cells showed nearly complete methylation of the CpG sites while DKO cells had minimal DNA methylation (**Figure 2A**). *DNAJB6* isoform expression was evaluated by isoform-specific qRT-PCR, and consistent with previous poly(A) sequencing data, DKO cells preferentially expressed the proximal poly(A) isoform when compared with HCT116 cells (**Figure 2B**). To test if isoform expression is dependent on DNA methylation, we treated HCT116 cells with the DNA demethylation agent, 5-aza-2’-deoxycytidine (DAC). DAC treatment resulted in the expected decrease in DNA methylation in HCT116 cells (**Figure 2A**) as well as an increase in proximal poly(A) isoform expression (**Figure 2B**). The shift in relative mRNA isoform abundance correlated with a similar trend in protein expression (**Figure 2C**). At baseline, HCT116 cells had both DNAJB6 proximal and distal protein expression while DKO cells had robust proximal protein expression but minimal distal protein expression. With DAC treatment, proximal protein increased in HCT116 cells, mirroring the increase in proximal poly(A) mRNA expression.

**Figure 2.**
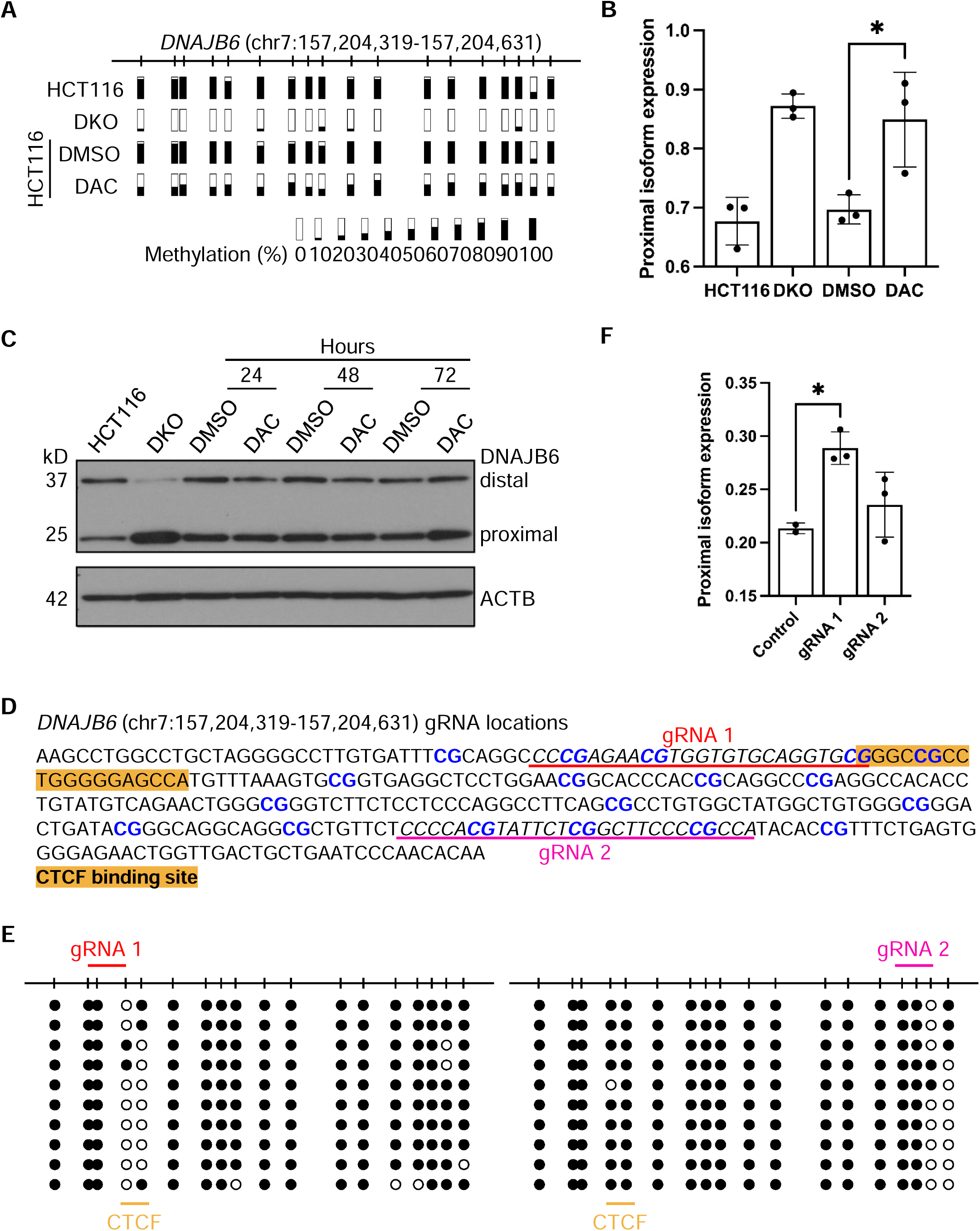
Methylation status of *DNAJB6* APA control region correlates with poly(A) isoform expression pattern. (**A**) Amplicon bisulfite sequencing of the putative APA control region in HCT116 and DKO cells as well as HCT116 cells treated with either DMSO (control) or DAC for 72 hours. Each bar represents a single CpG dinucleotide. The methylation level for at least 10 alleles is summarized for each sample. (**B**) *DNAJB6* proximal poly(A) isoform expression expressed as a fraction of the total *DNAJB6* gene expression in the same cells as in (A). The asterisk denotes statistical significance (p < 0.05). (**C**) Western blot for DNAJB6 protein isoform expression and beta actin (ACTB) loading control in the same cells as in (A). Earlier DAC treatment time points (24 hours and 48 hours) are also shown. (**D**) Design of guide RNAs (gRNAs) used in the dCas9-TET system for targeted DNA demethylation within the *DNAJB6* putative APA control region. Two gRNAs were used, and their respective position within the APA control region is underlined. The predicted CTCF binding site is highlighted in orange. (**E**) Amplicon bisulfite sequencing of the putative APA control region in HCT116 cells expressing the dCas9-TET system containing gRNA 1 (left panel) and gRNA 2 (right panel). Black circles represent methylated CpG, and white circles represent unmethylated CpG. Each row represents a single allele. (**F**) *DNAJB6* proximal poly(A) isoform expression expressed as a fraction of the total *DNAJB6* gene expression in the same cells as in (E). The control cells did not receive any gRNAs. The asterisk denotes statistical significance (p < 0.05).

While the above results supported that DNA methylation drives *DNAJB6* isoform expression, the genome-wide effects of DAC treatment cannot be ignored as possible confounding factors. Thus, we utilized a dCAS9-mediated epigenome-editing tool (21) to specifically demethylate CpG sites within the putative APA control region. Briefly, a fusion protein (dCAS-TET1-CD) between the catalytic domain of tet methylcytosine dioxygenase 1 (TET1; responsible for active DNA demethylation) and a dead CAS9 was targeted to the putative *DNAJB6* APA control region using two different guide RNA’s (**Figure 2D**). Co-expression of the guide RNA (gRNA) with dCAS-TET1-CD in HCT116 cells led to demethylation of specific CpG sites (**Figure 2E**). Isoform-specific RT-PCR revealed that cells expressing gRNA 1 significantly increased their proximal poly(A) isoform expression while cells expressing gRNA 2 did not shift isoform expression patterns significantly (**Figure 2F**). Interestingly, gRNA 1 targeted demethylation of CpG sites that overlap with the predicted CTCF binding motif while gRNA 2 targeted a downstream region that had less impact on isoform expression. Further manipulation of the extent of DNA demethylation within this region by using multiple different gRNAs tiling across this CpG island will help to identify more critical CpG sites in the future. Overall, these results suggest that *DNAJB6* isoform expression is indeed determined by the DNA methylation pattern of this CpG island.

### *DNAJB6* isoform expression and DNA methylation are regulated in response to heat shock

Next, we wondered whether any physiological stimuli could promote dynamic changes in *DNAJB6* isoform expression. We selected heat shock (HS) as a physiological stressor, given the known roles of DNAJB6 as a co-chaperone protein involved in maintaining proteostasis. We hypothesized that APA of *DNAJB6* might be dynamic under conditions where this protein has an important function (26). To test this, we isolated RNA, DNA, and protein from HCT116 cells at several time points following HS, to capture both the acute and recovery phases. Interestingly, we observed a shift in *DNAJB6* mRNA and protein expression towards the predominantly cytoplasmic proximal isoform immediately after HS, with the pattern persisting for 4 hours and recovering back to pre-HS levels at 24 hours (**Figure 3A** and **B**).

**Figure 3.**
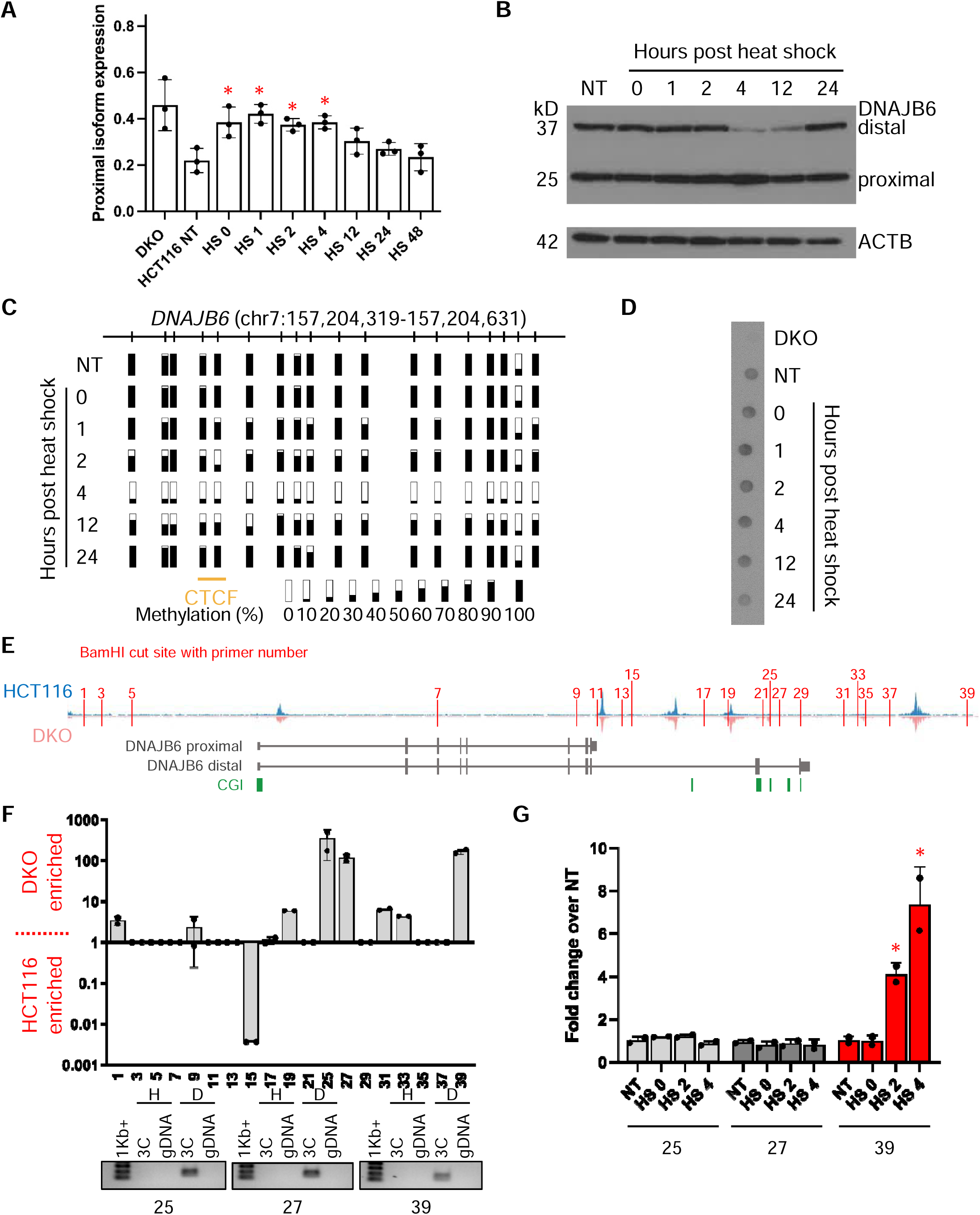
Heat shock induces DNAJB6 poly(A) isoform switching and targeted epigenetic remodelling. (**A**) *DNAJB6* proximal poly(A) isoform expression expressed as a fraction of the total *DNAJB6* gene expression in DKO, untreated HCT116 (NT), and post-heat shock HCT116 cells at the indicated time points (HS # where the number indicates hours post-heat shock). Asterisks identify heat shock (HS) samples that are statistically different from untreated (NT) HCT116 (p < 0.05). (**B**) Western blot for DNAJB6 protein isoform expression and beta actin (ACTB) loading control in the same HCT116 cells as in (A). (**C**) Amplicon bisulfite sequencing of the APA control region in same cells as in (B). Each bar represents a single CpG dinucleotide. The methylation level for at least 10 alleles is summarized for each sample. (**D**) Dot blot assay measuring the global 5-methylcytosine level in the same cells as in (A). (**E**) Schematic of the 3C assay fragments (defined by BamHI cut sites marked as red vertical lines) overlayed on top of the CTCF ChIP-seq data in HCT116 (blue) and DKO (pink) cells. The numbers correspond to the PCR primers used in the 3C assay and listed in Supplemental Table 3. (**F**) Fold enrichment of each 3C fragment amplification in DKO cells over HCT116 cells. Numbers correspond to PCR primers noted in (E). Realtime PCR products were resolved on 2% agarose gels for specificity and PCR product size confirmation. The images for the three most enriched fragments (primers 25, 27, and 39) in DKO cells were shown at the bottom. (**G**) Fold enrichment of 3C fragment amplification in post-heat shock HCT116 cells (HS # where the number indicates hours post-heat shock) over untreated HCT116 cells (NT) for primers 25 (light grey bars), 27 (dark grey bars), and 39 (red bars). The asterisk identifies HS 4 as statistically different from NT (adjusted p < 0.05).

To test whether DNA methylation was also affected, we preformed targeted bisulfite sequencing on the DNA isolated from these same cells. Strikingly, we observed dynamic changes in the DNA methylation pattern of the APA control region, with almost complete demethylation occurring by 4 hours post-HS and recovering to fully methylated at 24 hours post-HS (**Figure 3C**). This quick loss of DNA methylation indicates an active DNA demethylation mechanism, rather than passively losing DNA methylation through cell division. Furthermore, global DNA methylation levels remained mostly unchanged, as indicated by measuring total 5-methylcytosine content using a DNA dot blot (**Figure 3D**), suggesting that the dynamic DNA methylation remodelling is specifically targeted to the *DNAJB6* APA control region.

To test whether other cell stress-related mechanisms might be contributing to the *DNAJB6* APA isoform expression dynamic, we measured mRNA half-life of both *DNAJB6* isoforms following HS to determine whether differential rates of mRNA degradation might be contributing to the shift in isoform expression. We observed no significant difference in mRNA half-life between the distal and proximal isoforms following HS, with both isoforms remaining similarly stable (data not shown).

In our model of DNA methylation-regulated APA, preferential proximal poly(A) isoform expression is achieved by CTCF-mediated chromatin loop formation between the APA control region and distal CTCF binding sites (19). Leveraging chromatin conformation capture (3C) assay, we detected several distal genomic regions that interact with the *DNAJB6* APA control region as a function of differential DNA methylation (**Figure 3E** and **F**). Of note, chromatin contacts between the APA control region and BamHI fragments assayed by primers 25, 27, and 39 were the most enriched in DKO cells, identifying these as regions of interest to test in the HS time course. At 2- and 4-hours post-HS, we observed increased interaction between the APA control region and BamHI fragment assayed by primer 39 (**Figure 3G**). This fragment (chr7:157,222,183-157,233,363, GRCh37/hg19) is about 18 kb away in the 3’ direction from the APA control region, and it contains a clear CTCF binding site, as previously assayed by ChIP-seq (**Figure 3E**). The 3C results strongly suggested that a CTCF-mediated chromatin loop was formed upon DNA demethylation of the APA control region, consistent with our working model. It is worth noting that the 3C-based detection of chromatin loops is limited by user-defined regions; thus, future studies looking for global chromatin architecture changes during HSR will be highly important to expand our understanding of potential 3D genome restructuring as cells respond to stress. In one study, Hi-C analysis was utilized to assess global chromatin changes during acute HS and found that topologically associating domains and compartment structures stayed mostly unchanged in human and *Drosophila* (37). However, our findings at *DNAJB6* suggests that dynamic chromatin structure changes over time following HS may have been overlooked previously.

Collectively, these observations demonstrated that HS stress triggered a rapid change in *DNAJB6* poly(A) isoform expression pattern and invoked targeted DNA demethylation at the APA control region as part of the regulation. It is interesting to note that isoform switching preceded DNA demethylation in this time course, suggesting that additional signal transduction must be involved during the early phase of HSR to modulate APA and recruit DNA methylation modifiers for a sustained transcriptional countermeasure to this cellular stress.

### Identification of factors mediating DNA methylation-regulated APA during cell stress

As noted above, we speculated that additional factors participated in the HS-induced APA. The CTCF binding motif predicted in mediating DNA methylation-regulated APA at *DNAJB6* is only 19 bp in a 239 bp CpG island (**Figure 4A**), but the DNA methylation changes that occurred during HSR encompassed the entire CpG island. Therefore, we performed DNA pulldown followed by mass spectrometry to discover additional proteins involved in mediating the observed DNA methylation and chromatin structure changes. We used the entire CpG island as the DNA bait and performed the experiment with either methylated or unmethylated baits to enable identification of both DNA methylation-sensitive and insensitive binding proteins (**Figure 4B**). As a control, we probed the eluted proteins for CTCF and saw robust CTCF enrichment in the reaction containing the unmethylated bait (**Figure 4C**). After filtering out non-specific and low abundance proteins from replicate experiments, we identified 139 candidate binding proteins for the *DNAJB6* APA control region (**Supplemental Table 2**). A small subset of these proteins, including methyl-CpG binding domain protein 2 (MBD2) and E3 ubiquitin-protein ligase UHRF1 (UHRF1), showed DNA methylation-sensitive binding of the DNA baits, but the majority seemed to bind the DNA baits regardless of methylation status. To help prioritize for validation and functional studies, we annotated the protein list using GO terms and manually curated proteins with sequence-specific DNA binding (GO:0043565) for known functions in epigenetics, cell stress, and chromatin looping.

**Figure 4.**
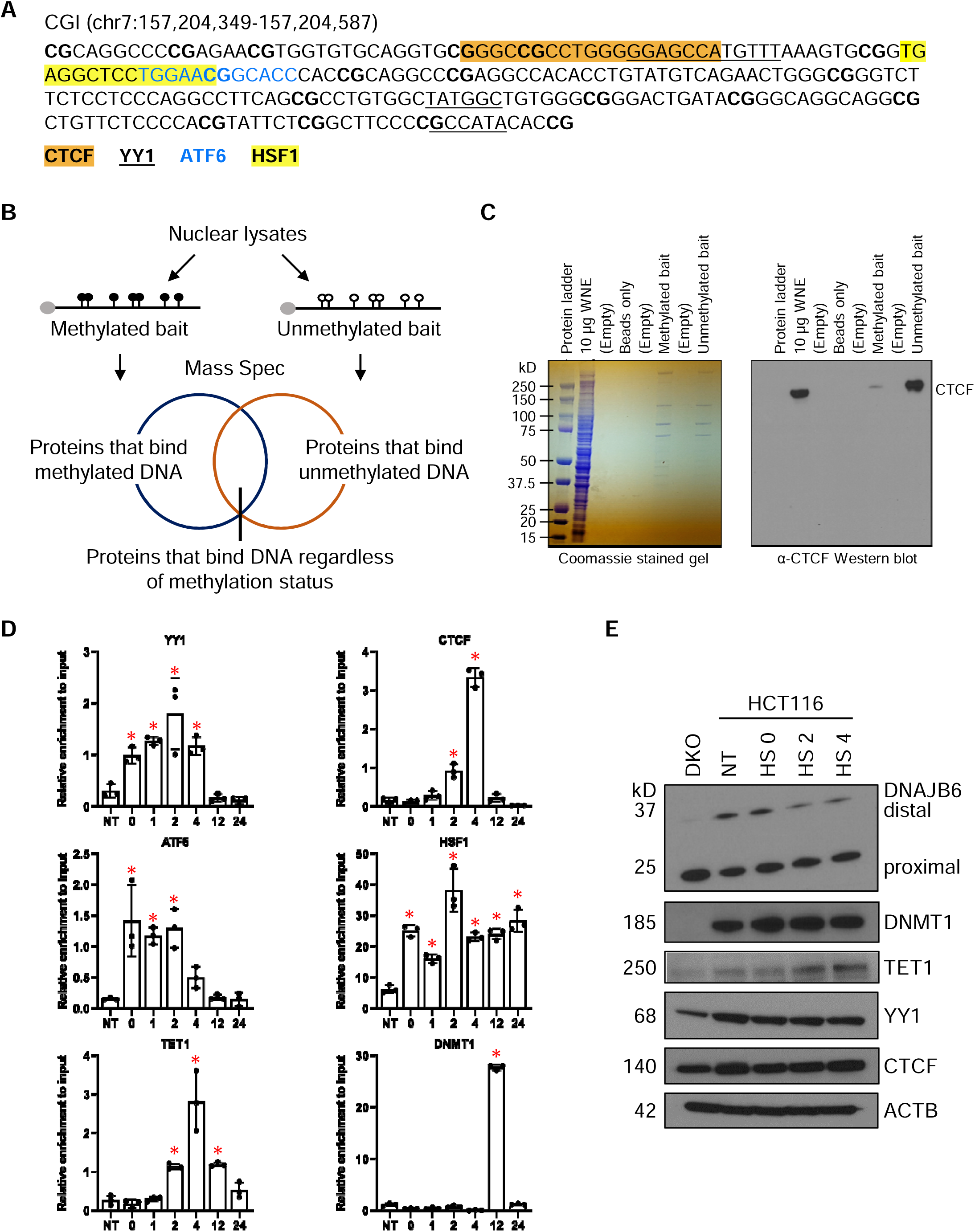
Transcription factors and DNA methylation modulators bind to the *DNAJB6* APA control region. (**A**) Annotation of the CpG island (CGI) sequence in the APA control region. Predicted binding sites for CTCF (highlighted in orange), YY1 (underlined), ATF6 (blue letters), and HSF1 (highlighted in yellow) are marked. (**B**) Workflow of DNA pulldown experiment. (**C**) Images of DNA pulldown cell lysates resolved on acrylamide gels and stained with Coomassie (left panel) and probed with anti-CTCF antibody (right panel). WNE, whole nuclear extract; Beads only, elute from reactions without nuclear extracts; Methylated bait, elute from reactions containing methylated bait; Unmethylated bait, elute from reactions containing unmethylated bait. (**D**) ChIP-qPCR in untreated HCT116 cells and post-heat shock (HS) HCT116 cells at the indicated time point. Numbers indicate hours post heat shock. Asterisks identify heat shock (HS) samples that are statistically different from untreated (NT) HCT116 (p < 0.05). (**E**) Western blot for DNAJB6 protein isoforms, DNMT1, TET1, YY1, CTCF, and ACTB loading control in DKO, untreated HCT116 (NT), and post-heat shock HCT116 cells at the indicated time points.

Based on the above criteria, we focused on Yin Yang 1 (YY1). YY1 is a constitutively expressed transcription factor known to contribute to chromatin loop formation, similarly to CTCF, in the context of promoter-enhancer interactions (38). In fact, co-binding and crosstalk between CTCF and YY1 have been reported across a variety of cell types and developmental stages, especially in the neural lineage (39-41). Sequence analysis of the APA control region identified three putative YY1 binding sites, one of which directly overlapped the CTCF motif (**Figure 4A**). Interestingly, this analysis also revealed potential binding of ATF6 and HSF1, both of which have well established roles in signal transduction during HSR and UPR (42-45), within this same genomic region. Because the DNA pulldown experiment was limited to untreated conditions, we wanted to explore in more detail the DNA binding dynamics of CTCF, YY1, ATF6, and HSF1, as well as DNA methylation regulators, DNA methyltransferase 1 (DNMT1) and tet methylcytosine dioxygenase 1 (TET1). To this end, we preformed chromatin immunoprecipitation on HCT116 cells following HS treatment and observed immediate binding of ATF6, HSF1, and YY1 at the APA control region (0-hour time point in **Figure 4D**), concurrent with the increase in proximal poly(A) isoform expression (**Figure 4A**). However, there appeared to be slight differences in the binding dynamic of these three transcription factors; HSF1 localization persisted throughout the entire time course, ATF6 binding tapered off by 4-hour post-HS, while YY1 binding peaked at 2-hour post-HS. This observed pattern likely resulted from differences in how these transcription factors operate during stress response; HSF1 activation is achieved through homotrimer formation upon release from molecular chaperone proteins (46,47) and via direct thermo-sensing (48), soluble ATF6 is released from ER by proteolysis so that it can translocate into the nucleus (49), and finally, ATF6 and YY1 co-activate transcription at ER stress element (ERSE)-containing promoters during UPR (50).

Furthermore, we detected TET1 recruitment to the APA control region starting at 2- hour post-HS and peaking at 4-hour, coinciding with the near complete demethylation of this region seen by targeted bisulfite sequencing (**Figure 3C**). TET1 is one of three enzymes responsible for active DNA demethylation in mammalian cells, converting 5-methylcytosines to 5-hydroxy-, 5-carboxyl-, and 5-formyl-cytosines in a stepwise fashion (51-53). Therefore, its localization to the APA control region provided a molecular explanation for the rapid DNA demethylation observed at this region. Tracking nicely with the methylation status of its binding site (**Figure 3C**), CTCF was also detected here starting at 2-hour post-HS and disappearing by the 12-hour time point (**Figure 4D**). Finally, as the cells recovered from the heat stress and restored *DNAJB6* proximal poly(A) isoform expression back to homeostatic level, DNMT1 was found at this region to facilitate DNA re-methylation (**Figure 4D**). Interestingly, only TET1 protein level seemed to have increased during this time course (**Figure 4E**). While we were not aware of published reports on HS inducing TET1 transcription, scanning the TET1 promoter sequence identified a HSF1 motif (-264 bp from transcription start site) and two ATF6 motifs (-269 bp and +43 bp from transcription start site), raising the possibility that TET1 expression could be stimulated in response to cellular stress.

### Discussions

Altogether, our data suggested that the *DNAJB6* APA control region contains sequence elements that allow for binding and integration of stress-sensing transcription factors (HSF1 and ATF6) to quickly increase *DNAJB6* proximal poly(A) isoform expression during HSR and produce the corresponding cytoplasmic protein isoform capable of suppressing protein aggregate formation. Transcriptional co-regulators, such as YY1, can also bind to this same region to facilitate isoform switching. TET1 is then recruited to rapidly demethylate the APA control region, which enables CTCF binding and the ensuing chromatin loop formation. Based on our prior work, this CTCF-mediated chromatin looping can further enforce proximal poly(A) mRNA expression, thus ensuring reliable proximal protein production to support cell needs. As the cells recover from HS, the transcription factors terminate their tenure at the APA control region, DNMT1 is recruited to remethylate the DNA, and the *DNAJB6* APA rheostat is reset.

Combinatorial transcription factor binding is thought to contribute to the evolutionary stabilization of DNA regulatory regions; therefore, this co-occurrence of YY1, ATF6, HSF1, and CTCF binding, overlaying the dynamic DNA methylation scenery, highlights the *DNAJB6* APA control region as an important and tuneable regulatory element of gene expression, akin to a gene promotor. Moreover, the 3D chromatin loop formation (**Figure 3F** and **3G**) and the enrichment of histone H3 lysine 27 acetylation (H3K27Ac in **Figure 1B**) and serine 5 phosphorylated RNA polymerase II (Pol2Ser5 in **Figure 1B**) at this same region strikingly resemble classical enhancer-promoter interactions that regulate transcription activity of single poly(A) site genes. Recently, enhancers have been shown to regulate APA in addition to modulating the total transcriptional output of multi-UTR genes (54). In this study, perturbations of YY1, CTCF, and cohesin complex components altered the cleavage and polyadenylation activity of a reporter system for *PTEN* APA. Thus, we surmise that the distal interacting BamHI fragment detected in the 3C experiment (**Figure 3G**) potentially contains an enhancer element that further contributes to the fine-tuning of *DNAJB6* APA. While there is robust literature on stress-induced gene activation and silencing, our findings here represent a novel example of stress-induced transcriptome modulation, where transcription factors, DNA methylation, and chromatin architecture converge to achieve a necessary poly(A) isoform expression pattern.

Our observations also raised several important questions that warrant further investigation. First, how universal is this mechanism during different types of cellular stress response? We speculate that stressors that elicit overlapping signal transduction pathways might induce similar changes in chromatin and isoform expression in *DNAJB6* and other potential targets like it. For instance, drugs that cause ER stress, such as tunicamycin and thapsigargin, are known to activate ATF6 and can potentially trigger similar APA and chromatin changes described in this study. Second, what are the genes subjected to this type of APA regulation? While we came upon *DNAJB6* through our prior work investigating the function of gene body DNA methylation, heat shock, and perhaps other cellular stress conditions, offers an excellent context to identify physiologically relevant APA globally using genomic approaches. Third, the swift DNA methylation changes and the sequential binding of transcription factors preceding methylation changes in the HS time course presents an exquisite opportunity for investigating stress-induced DNA methylome remodelling. An obvious question is whether YY1 facilitated the recruitment of TET1. It has been reported the YY1 has DNA methylation sensitivity, but the results from the DNA pulldown experiment indicated that it can bind regardless of DNA methylation, in contrast to CTCF. It is therefore tempting to speculate the necessity of YY1 binding in facilitating the DNA de-methylation machinery and subsequent CTCF binding. Further studies expanding upon the global chromatin occupancy of these factors under dynamic conditions will be important for unravelling this mechanism of gene regulation further. Finally, it is worth noting that *DNAJB6* poly(A) isoforms can also be alternatively spliced to generate different 3’ coding regions and 3’ UTR. Alternative splicing and APA are often coupled during mRNA processing. However, delineating the relative contribution of alternative splicing and APA in regulating isoform expressions or how each process impacts the other is beyond the scope of this current study but will be highly interesting to investigate in the future.

## DATA AVAILABILITY

ChIP-seq data for CTCF, SMC1, RAD21, Pol2Ser5, and H3K27Ac are available under accession number GEO:GSE131606 (19).

## SUPPLEMENTARY DATA

Supplementary Data are available at NAR online.

## ACKNOWLEDGEMENT

We acknowledge the Proteomics and Metabolomics Core at the Lerner Research Institute, Cleveland Clinic, for their expert assistance in performing the mass spectrometry analyses in the DNA pulldown experiment.

---

**Supplemental Table 1.** GO analysis of 546 candidates of DNA methylation-regulated APA.

**Supplemental Table 2.** Annotation of 139 candidates from DNA pulldown experiments.

**Supplemental Table 3.** PCR primers used in this study.

