## Supplementary material for "Heat shock induces alternative polyadenylation through dynamic DNA methylation-regulated chromatin looping": Table S3

**Supplemental Table 3. Primers used in this study**

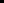

| **Primer name** | **Sequence** | **Amplicon size (bp)** |
| --- | --- | --- |
| **qRT-PCR** | | |
| DNAJB6 distal-FW | GAACTCATGTTTCAGTTCGCG | 145 |
| DNAJB6 distal-BW | GGAATTCTACTCCGTGCTCAA |  |
| DNAJB6 proximal-FW | AGCTCATCGGAGCCTCTATTT | 160 |
| DNAJB6 proximal-BW | TACACTAAAGAGTCAAGACTCAC |  |
| **DNA pulldown probe** | | |
| DNAJB6-fwd | GTCTTTGCAAGGTGCAGGAT | 409 |
| DNAJB6-rev | TGGCCTCTTTGCTCTAAGGA |  |
| **ChIP qPCR** | | |
| DNAJB6 set 1-F | TGACTGCTGAATCCCAACAC | 106 |
| DNAJB6 set 1-R | AGAGGAAAAGCCGCAAAACG |  |
| **Bisulfite sequencing** | | |
| DNAJB6-BSF-18-F1 | AAGTTTGGTTTGTTAGGGGTTTT | 313 |
| DNAJB6-BSF-18-R1 | AAAATTCAACAATCAACCAATTCTC |  |
| **Chromosome conformation capture (3C)** | | |
| 3C-DBAM-22 (anchor) | CCTCACCGCACTTTAAACAT | N/A |
| 3C-DBAM-1 | CCAGCCTGTAATCCCAGCTA | 265 |
| 3C-DBAM-3 | AAGACTCGTCACCCATGTCC | 316 |
| 3C-DBAM-5 | CCCAGTGACCTCACTCACCT | 269 |
| 3C-DBAM-7 | TGTCGGATGGTGAGTGACAG | 313 |
| 3C-DBAM-9 | AGCAGGCCCAGCTAATTTTT | 325 |
| 3C-DBAM-11 | ACAGGAGAGGGAAGAAACCA | 315 |
| 3C-DBAM-13 | AGCTGCCTGGAGTGTAGAGC | 332 |
| 3C-DBAM-15 | GGTCTCAGCCCAAGTCTTCC | 263 |
| 3C-DBAM-17 | GATGGGGTCTCCCTATGTTG | 324 |
| 3C-DBAM-19 | ACACCTGCCTTGCTCTCTGT | 263 |
| 3C-DBAM-21 | AGCAGGACATGAAAAGAAGG | 395 |
| 3C-DBAM-25 | CATTTGCTGCTGTGTGGAGT | 323 |
| 3C-DBAM-27 | GACAGACGTTTGGGACTTGG | 319 |
| 3C-DBAM-29 | TTTCCTTCCTGGGCGTCT | 336 |
| 3C-DBAM-31 | GAAAAGAAGCCTTCCCGAGT | 315 |
| 3C-DBAM-33 | ACCTGCAGCTTCAGATCCAC | 310 |
| 3C-DBAM-35 | GATGTTTGTGCTCACGTGCT | 297 |
| 3C-DBAM-37 | ACACTGAGCAGGGTGAGCTT | 279 |
| 3C-DBAM-39 | TGGGATTCTTGGTTCTGCTC | 310 |
